# Neural Entrainment to Musical Pulse In Naturalistic Music Is Preserved In Aging: Implications for Music-Based Interventions

**DOI:** 10.1101/2022.11.05.515314

**Authors:** Parker Tichko, Nicole Page, Ji Chul Kim, Edward Large, Psyche Loui

## Abstract

Neural entrainment to musical rhythm is thought to underlie the perception and production of music. In aging populations, the strength of neural entrainment to rhythm has been found to be attenuated, particularly during attentive listening to auditory streams. However, previous studies on neural entrainment to rhythm and aging have often employed artificial auditory rhythms or limited pieces of recorded, naturalistic music, failing to account for the diversity of rhythmic structures found in natural music. As part of larger project assessing a novel music-based intervention for healthy aging, we investigated neural entrainment to musical rhythms in the electroencephalogram (EEG) while participants listened to self-selected musical recordings across a sample of younger and older adults. We specifically measured neural entrainment to the level of musical pulse—quantified here as the phase-locking value (PLV)—after normalizing the PLVs to each musical recording’s detected pulse frequency. As predicted, we observed strong neural phase-locking to musical pulse, and to the sub-harmonic and harmonic levels of musical meter. Overall, PLVs were not significantly different between older and younger adults. This preserved neural entrainment to musical pulse and rhythm could support the design of music-based interventions that aim to modulate endogenous brain activity via self-selected music for healthy cognitive aging.

## 1. Introduction

Music-based interventions (MBIs), such as receptive MBI (i.e., interventions that involve listening to music), have become increasingly of interest for improving well-being across the lifespan (Cheever et al., 2018; Global Council on Brain Health, 2020). Despite the growing inclusion of MBIs into healthcare protocols, meta-analyses suggest that they often produce variable and inconsistent effects on clinical and health-related outcomes (Mammarella et al., 2007; Sousa et al., 2020; van der Steen et al., 2018; Vasionytė & Madison, 2013; Vink & Hanser, 2018). Such variability in the efficacy of music-based interventions may arise, in part, from the diversity of protocols that underlie MBIs (e.g., self-selected vs. clinician-selected music), the heterogeneity of clinical populations that are targeted by MBIs, and individual differences in the sensitivity to musical features that constitute the intervention (e.g., rhythm, melody, motor-movement, and social interactions during musical experiences) (Loui, 2020; Sousa et al., 2020; Vink & Hanser, 2018). While research has identified key neural networks that contribute to music processing (Koelsch, 2014; Loui & Przysinda, 2017), little is known about the underlying neurobiological mechanisms that are *specifically* engaged by MBIs (Quinci et al., (2022); Wang et al., (2020)) and how aging affects neural responses to musical structure (e.g., rhythm, melody, harmony) (Sauvé et al., 2019; Sutcliffe et al., 2020). Yet, understanding how MBIs engage the nervous system and the impact of aging on the neural processing of music has important implications for designing and implementing MBIs; understanding naturalistic music-listening and -making on brain function, cognitive health, and well-being; and explaining individual outcomes following the intervention (Ferreri et al., 2019; Sutcliffe et al., 2020; Tichko et al., 2020).

Music engages multiple neural systems that subserve sensorimotor functions, executive control, reward processing, and vestibular function (Koelsch, 2014; Loui & Przysinda, 2017; Vuust et al., 2022). Musical experiences involve a listener’s idiosyncratic musical knowledge, autobiographical memories, affective state, and subjective interpretation of a composer’s and performer’s musical intentions (Alluri et al., 2017; Aydogan et al., 2018; Janata, 2009; Juslin & Laukka, 2004; Sloboda et al., 2001). This inherent subjectivity of musical experiences may explain why self-selected music produces enhanced brain responses to music (Blood & Zatorre, 2001; Loui, 2020; Quinci et al., 2022; Salimpoor et al., 2013). Neuro-imaging research, for example, has found that listening to self-selected music and music perceived as pleasurable increases activation in and connectivity between auditory and reward systems (Blood & Zatorre, 2001; Salimpoor et al., 2013) (Ferreri et al., 2019; Gold et al., 2019). Furthermore, music that is more familiar and selected by the listener is especially effective at engaging multiple brain areas (Pereira et al., 2011): one study showed increased functional connectivity between auditory and reward systems when participants listened to self-selected music, with effects increasing after a two-month MBI (Quinci et al., 2022). These findings may be linked to the observation that music-based interventions that feature self-selected music yield better clinical outcomes (Leggieri et al., 2019), such as in anxiety reduction and improvements in task performance and enjoyment (Cassidy & Macdonald, 2009).

In addition to engaging auditory, motor, and reward systems, music also engages neural oscillations—patterns of rhythmic activity arising from excitatory-inhibitory neuronal interactions. During music-listening, endogenous oscillations in auditory-motor systems (Arnal et al., 2015; Fujioka et al., 2012; Nozaradan et al., 2012) their activity to rhythmic timescales in music (Fujioka et al., 2012, 2015; Harding et al., 2019; Nozaradan et al., 2012; Stefanics et al., 2010; Will & Berg, 2007; Woods et al., 2021). For example, musical pulse occurs naturally within a frequency range of 0.5-4 Hz, with a prominent frequency generally centered around 2 Hz (Ding et al., 2017; Large et al., 2015). Musical meter includes the pulse frequency, sub-divisions of the pulse level that occur between 4-8 Hz, and slower beats that group pulse cycles (< 2 Hz). These pulse and metrical frequencies overlap with delta (e.g., 0.5-4 Hz) and theta (e.g., 4–8 Hz) bands of endogenous activity generated by the brain (Large et al., 2015). Electroencephalographic (EEG) and magnetoencephalographic (MEG) recordings of brain activity taken during music-listening have observed the entrainment of delta and theta responses to the rhythmic structure of music, resulting in increased power, phase-locked responses, and mode-locked responses at frequencies related to pulse and meter (Doelling & Poeppel, 2015; Fujioka et al., 2012; Nozaradan et al., 2011; Stefanics et al., 2010; Will & Berg, 2007; Woods et al., 2021). Such phase- and mode-locked responses have been recorded to auditory stimuli falling within the range of human rhythm (Harding et al., 2019; Tal et al., 2017; Vanden Bosch der Nederlanden et al., 2020; Woods et al., 2021) and pitch (Lerud et al., 2014; Skoe et al., 2013; Skoe & Kraus, 2010; Tichko & Skoe, 2017) perception, suggesting that neural entrainment may be a general, dynamical property underlying brain function (Tognoli & Kelso, 2009).

While music has been shown to entrain neural activity, impairments in neural entrainment to musical features are associated with aging (Alain et al., 2014; Bones & Plack, 2015; Henry et al., 2017; Sauvé et al., 2019; Sutcliffe et al., 2020; Zendel & Alain, 2014). Relatedly, several neurodegenerative disorders, such as Mild Cognitive Impairment and Alzheimer’s Disease, are associated with disrupted neural activity across the same frequency bands that are driven by music (e.g., delta, theta) (Goodman et al., 2018; Güntekin & Başar, 2016; Hata et al., 2016; Koenig et al., 2005). These findings have motivated the development of non-invasive interventions that aim to entrain aberrant brain activity with rhythmic stimulation to promote healthy cognitive aging (Reinhart & Nguyen, 2019) and slow disease progression (Iaccarino et al., 2016; Martorell et al., 2019).

The reviewed work suggests that entraining brain activity through music could be an effective strategy for treating neurodegenerative disorders with MBIs. However, research on neural entrainment to music has typically used non-ecological musical stimuli (e.g., with artificial auditory rhythms (e.g., amplitude-modulated tones) or short excerpts of natural music that represent a few prominent pulse frequencies (e.g., (Doelling & Poeppel, 2015; Harding et al., 2019; Henry et al., 2017; Nozaradan et al., 2011; Will & Berg, 2007)) that do not represent the rich musical experiences that often constitute MBIs. In contrast to the design of these studies, MBIs, especially those that allow participants to select their own musical materials, employ a range of natural music with rhythmic structures that reflect varying pulse and metrical frequencies. Even within a single piece of music, pulse and tempo fluctuations can occur to communicate expressive intent and large-scale musical structure (Ashley, 2002; Chapin et al., 2010; Istók et al., 2013). This inherent variability in rhythmic content and fluctuations in tempi pose a methodological challenge for analyzing neural entrainment to music under listening conditions that are more representative of current MBIs (e.g., across musical stimuli that are designed to feature different pulse frequencies or for musical stimuli that are selected by a listener and cannot be experimentally controlled).

Neurodynamical models of musical rhythm, such as gradient-frequency neural networks (Kim & Large, 2015, 2019, 2021; Lambert et al., 2016; Large et al., 2010, 2015; Velasco & Large, 2011) which simulate the entrainment of neural ensembles to musical rhythm, have successfully explained and predicted neural, behavioral, and psychological responses to music in human listeners (Large et al., 2015; Tal et al., 2017; Tichko et al., 2021; Tichko & Large, 2019). In addition to accounting for human responses to music, these models can also be used for signal-processing of biological signals (Kaplan & Chew, 2019) and for music-feature analysis, including beat-finding, tempo-tracking, and chord estimation (Kim, 2017; Lambert et al., 2016; Large et al., 2015). This suggests that neurodynamical models of music could inform the analysis of neural entrainment to more naturalistic musical stimuli by estimating pulse and meter-related frequencies that are likely to be perceived by human listeners.

In the current study, we investigated younger and older adults’ neural entrainment to musical pulse during a period of non-invasive, audiovisual stimulation that featured self-selected music and music-synchronizing lights. We used a modified version of a neurodynamical model of human pulse perception (Large et al., 2015) to estimate the perceived pulse and metrical frequencies for each musical recording, and then measured neural entrainment to musical pulse, after accounting for each musical recording’s unique pulse and metrical frequencies. Motivated by the reviewed work, we predicted that we would observe enhanced neural entrainment to the level of musical pulse in self-selected music, relative to non-rhythmic levels (Doelling & Poeppel, 2015; Fiveash et al., 2020; Vanden Bosch der Nederlanden et al., 2020). Secondly, we predicted that neural responses to the pulse would be relatively stronger at fronto-central electrodes, reflecting synchronized neural activity to music arising from the auditory system (Fiveash et al., 2020; Vanden Bosch der Nederlanden et al., 2020; Woods et al., 2021). Finally, we expected younger adults to exhibit stronger neural entrainment to musical pulse, compared to older adults, given that degraded neural responses to sound are often associated with aging (Henry et al., 2017; Sauvé et al., 2019).

## 2. Materials and Methods

### 2.1. Participants

16 young adults (Mean age = 19.81 years, range = 18–22 years; 7 Males, 9 Females) and 16 older adults (Mean age = 70.94 years, range = 55–81 years; 5 Males, 11 females) were recruited for a behavioral and electrophysiology (EEG) study in the Music Imaging and Neural Dynamics (MIND) Lab at Northeastern University. Across the younger and older adults, 94% of participants reported a predominant handedness of right (30 right handedness, 2 left handedness) and 93% reported a first language of English (28 English, 1 Fulani, 1 Norwegian, 1 Mandarin, 1 German). The young adults participated in return for course credit at Northeastern University. The older adults were compensated at $20 per hour for their participation. The study was approved by the IRB of Northeastern University (IRB #19-03-20). After informed consent, participants completed a battery of behavioral tasks, as described in Behavioral Battery below.

### 2.2. Procedure

As part of a larger project on the development of a novel music-based intervention for aging, participants underwent a single session of audiovisual stimulation, which consisted of listening to self-selected music while viewing music-synchronized lights. Next, participants completed a visual working-memory task to assess cognitive functioning (Reinhart & Nguyen, 2019). Results on this working-memory task will be reported in a separate manuscript.

### 2.3. Behavioral Battery

Prior to the audiovisual stimulation, participants completed a behavioral battery to assess musical reward and sensitivity (Barcelona Music Reward Questionnaire (BMRQ)) (Mas-Herrero et al., 2013), musical sophistication (Goldsmiths Musical Sophistication Index (Gold-MSI)) (Müllensiefen et al., 2014), melodic contour perception (Montreal Battery of Evaluation of Amusia (MBEA)) (Peretz et al., 2003). Participants also self-reported basic demographic information, age, sex, medical history (e.g., hearing ability), length and nature of previous musical training, native languages, and handedness.

### 2.4. Audiovisual Stimulation

Prior to their in-lab session, participants selected six individual musical recordings for a period of naturalistic music listening while viewing light-emitting diode (LED) lights that synchronized to the pulse frequency of the music. Musical recordings were presented to the participants via a Macbook Air running a custom experiment script in PsychoPy Version 3.2.3 (Peirce et al., 2019) and Sennheiser CX 300-II earbuds. Music-synchronized lighting effects were created using a Synchrony™ device (Oscilloscape LLC), that analyzes the rhythmic structure of music in real-time and flashes LED lights synchronized to the music’s pulse. The display pattern on Synchrony™ was set to “color pulse,” and the color palette was set to third color palette (e.g., a set of cool colors). With these settings, the device softly pulses LED lights at the rate of musical pulse using a color palette of cool colors (e.g., blues, purples, whites) that change at the rate of the measure-level. For the younger adults, the visual stimulation was presented via a WS-2811 LED strip that was affixed to a table located over the legs of the participants in a seated position. During the audiovisual stimulation, younger participants were instructed to remain motionless, listen to the music, and engage with the lights by foveating directly at the LED strip. For the older adults, the visual stimulation was presented on a circular LED frame consisting of a WS-2811 LED strip, a circular metal frame with a 21” diameter, and a fixation cross located at the center of the frame, such that the LED strip was positioned at a 20° visual angle from the observer along the perimeter of the circular frame. This circular display was chosen to be visible to the participant through their peripheral vision, which is more sensitive to small changes in brightness in dim light situations due to the richness of rod cells in the retina at 10-20° from foveation (Purves et al, 2019). During the audiovisual stimulation, older participants were instructed to remain motionless, listen to the music, and engage with the lights by foveating on the fixation cross.

### 2.5. EEG Recording

Participants’ EEG was collected using a 64-channel BrainVision system, arranged according to the international 10-20 standard, and PyCorder software. EEG was recorded using an online reference of Fp1 and a 5000-Hz sampling rate. EEG time series were recorded directly to disk in the BrainVision format with trigger events that denoted the beginning and end of each musical recording. Impedances were kept <30 kOhm.

#### Analysis

### 2.5. EEG Preprocessing

The raw electroencephalogram was preprocessed using custom MATLAB (R2019a,b) routines and the EEGLab library (Delorme & Makeig, 2004) for MATLAB (versions 2020.1 & 2021.1). An initial epoch of the EEG data was created to remove superfluous activity unrelated to the audiovisual stimulation, i.e., activity that began 1 second prior to the start of the first musical recording and 1 second following the completion of the final musical recording. After this initial epoching, channel data were down-sampled from a 5000-Hz to 1000-Hz sampling rate. EEG channels were, then, re-referenced to bi-lateral mastoids at electrodes TP9 and TP10. To remove slow drifts and high-frequency activity unrelated to the audiovisual stimulation, EEG data were high-pass filtered at 1 Hz and low-pass filtered at 55 Hz using EEGLab’s Hamming windowed sinc finite impulse response (FIR) filter (*eegfiltnew*). Residual 60-Hz line noise was removed using CleanLine’s multi-taper filter with a threshold (i.e., p-value) set to 0.05. After filtering, bad channels were identified and removed using a semi-automated procedure: bad channels were automatically rejected using EEGLab’s joint-probability algorithm using a normalized threshold of 5 standard deviations. Remaining channels were then visualized, and additional noisy channels were removed manually after inspection by several trained research assistants (average number of channels removed per participant = 3.45, SD = 3.10). Following the removal of bad channels, non-linearities in the electroencephalogram were corrected using the Artifact Subspace Reconstruction (ASR) algorithm (Mullen et al., 2015) with a standard deviation threshold set to 20 and k-window set to 0.25–parameters which have been shown to correct artifact-driven non-linearities in EEG data, while preserving brain-related activity (Chang et al., 2020). Finally, EEG source decomposition was conducted, using independent components analysis, to remove components that reflected eye and cranial-muscle artifacts. After decomposition, the independent components (ICs) were subsequently classified using ICALabel, a pre-trained, machine-learning classifier that computes probabilities for multiple source classes (Pion-Tonachini et al., 2019). ICs classified as an eye or muscle source with a 90% probability were rejected automatically. Remaining ICs were manually inspected by trained research assistants using EEGLAB’s extended IC properties (e.g., IC power spectra, topographies, and time series) and removed if they contained extensive eye, muscle, or line-noise artifacts (average number of ICs removed per participant = 5.94, SD = 3.82). After removal of ICs, spherical interpolation was used to interpolate previously rejected channels. Finally, the full pre-processed EEG time series was epoched into six data sets per participant, each data set reflecting one of the participant’s self-selected musical recordings, to conduct recording-level analyses.

### 2.6. Music-Feature Analysis: Estimating Pulse Frequencies and Identifying Stable Epochs

Before estimating neural entrainment to musical rhythm, a music-feature analysis of each musical recording was conducted to identify a specific epoch that contained a stable pulse frequency. Identifying a single epoch for each musical recording with a stable pulse frequency for our analysis allowed us to obtain a more accurate estimate of neural entrainment at the pulse frequency by controlling for large tempo changes. Before the analysis of neural entrainment to musical rhythm, a music-feature analysis was conducted to identify an epoch of each musical recording that contained a stable pulse frequency. First, a modified version of the oscillator network model described in Large et al. (2015) was used to estimate pulse and metrical frequencies perceived by human listeners. The audio signal of each musical recording was processed with a middle-ear filter (Zilany & Bruce, 2006), and then complex-domain onset detection (Bello et al., 2004) was applied to derive an onset signal containing pulses triggered by the onset (i.e., attack) of individual musical events. The oscillator network was driven by the onset signal, and metrical frequencies (e.g., pulse, harmonic and/or subharmonic) were identified from peaks in the oscillator amplitudes. Stable epochs were identified using the same algorithm. The algorithm identified 1) the most salient frequency in the delta range (< =3.5Hz), 2) the most salient harmonic in the theta range (≥ 3.5Hz), and 3) the most salient subharmonic of the pulse. It then identified the first interval of at least 2 minutes in which all the frequencies remained stable within a few percent. The neural-entrainment analyses that follow were conducted over musical-recording epochs identified by our algorithmic method to have a consistent pulse frequency.

### 2.7. Neural Entrainment to Rhythm: Phase-Locking Values

Neural entrainment was operationalized as the phase-locking value (PLV) (Cohen, 2014; Chapter 19) across pulse- and meter-related frequencies—specifically 0.25-5 Hz — between the amplitude envelope of each musical recording and preprocessed EEG data. The amplitude envelope of each musical recording was estimated using the method by Lalor & Foxe, (2010), as implemented in the Multivariate Temporal Response Function (mTRF) toolbox (Crosse et al., 2016) using a sample rate of 1000 Hz. To compute PLVs, first, a complex wavelet transform of the amplitude envelope of the musical recording and each channel of preprocessed EEG data was conducted using logarithmically spaced complex Morlet wavelets from 0.25-5 Hz, with 15 bins per octave. The number of cycles per wavelet was determined algorithmically by doubling the center frequency the Morlet wavelet and rounding to the nearest integer. This produced a total of 65 Morlet wavelets with an increasing number of cycles (range of 3–8 cycles) and center frequencies that spanned the range of musical rhythm (Large et al., 2015). The amplitude envelopes of the musical recordings and the EEG-channel data were, then, convolved with the complex Morlet wavelets. Following the convolutions, phase angles were extracted from the complex-numbered time-series to estimate the instantaneous phase of the amplitude envelope of the musical recordings and EEG-channel data for each complex Morlet wavelet. From the phase-angle time-series, neural entrainment to music was quantified as the phase-locking value (PLV; Equation 1) for each EEG channel and each complex Morlet wavelet (Cohen, 2014; Chapter 19):

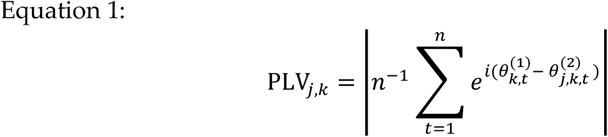

Here, PLV_j,k_ is the phase-locking value (PLV) of the jth EEG channel for the kth complex Morlet wavelet. 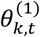 and 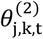 correspond to the instantaneous phase angles of the amplitude envelope of the music recording and EEG data, respectively; *t* corresponds to time point, *t*, in discrete time, e is Euler’s number, and *i* is the unit imaginary number. The PLV is a scalar value, (0 ≤ PLV ≤ 1), defined as the magnitude of the mean resultant vector calculated from the distribution of EEG-music relative phases. Relatively higher PLVs indicate stronger phase-locking between EEG signals and the amplitude envelope of the musical recording.

### 2.8. Pulse Normalization of Phase-Locking Values

Because participants selected musical recordings featured different pulse frequencies, we normalized the PLVs for each musical recording to a pulse frequency of 2 Hz prior to second-level analyses (i.e., before averaging PLVs across participants, electrodes, and musical recordings). Normalizing the PLVs to the same pulse frequency allowed us to investigate neural entrainment to musical pulse at the aggregate level in our groups of younger and older adults. For instance, if our music-feature analysis of the musical recording detected a prominent pulse frequency of 2.5 Hz, the phase-locking values would consequently be shifted in the frequency domain from 2.5 Hz to 2. Thus, in this analysis, the dimensionless unit of 2 corresponds to the pulse frequency (henceforth called the “pulse level”), 1 corresponds to a normalized subharmonic frequency (henceforth called the “subharmonic level”), and 4 corresponds to a normalized harmonic frequency (henceforth called the “harmonic level”). Motivated by previous analyses on the neural entrainment to auditory rhythms (Fiveash et al., 2020), we also selected a level between the pulse and harmonic level (dimensionless unit 3, henceforth called “off-pulse level”), to investigate whether neural entrainment was stronger at the predicted pulse level, relative to a neighboring, non-pulse level. Finally, for exploratory analyses of the sub-harmonic and harmonic levels, and we also defined an “off-subharmonic level” (unit 1.5) and an “off-harmonic level” (unit 5).

To test our *a priori* predictions (e.g., neural entrainment would emerge at the pulse level, neural entrainment would be stronger in younger adults) at the group-level, PLVs were averaged across each participants’ musical recordings and all EEG electrodes, yielding one grand-averaged PLV for each participant. For the grand-averaged PLVs, 95% confidence intervals were bootstrapped for each age group using with boot library for R (Canty & Ripley, 2021) with the normal approximation set to 10,000 samples (Davison & Hinkley, 1997). In addition to calculating grand-averaged PLVs, we also explored whether PLVs were strongest at fronto-central electrodes, consistent with an auditory response (Hall, 1992), and differed across clusters of EEG channels. Nine electrode clusters, consisting of 6 electrodes each, used previously in auditory-related EEG research (Riha et al., 2020) were selected for this analysis. Table 1 presents the 9 electrode clusters and their 6 constituent EEG channels.

**Table 1.** Nine electrode clusters, defined from previous auditory-related research, were used to investigate channel-related differences in neural entrainment to rhythm.

| Electrode Cluster | Electrodes |
| --- | --- |
| Left Frontal (LF) | AF7, AF3, F7, F5, F3, F1 |
| Right Frontal (RF) | AF4, AF8, F2, F4, F6, F8 |
| Left Central (LC) | FT7, FC5, FC3, T7, C5, C3 |
| Midline Central (MC) | FC1, FCz, FC2, C1, Cz, C2 |
| Right Central (RC) | FC4, FC6, FC8, C4, C6, T8 |
| Left Parietal (LP) | TP7, CP5, CP3, P7, P5, P3 |
| Midline Parietal (MP) | CP1, CPZ, CP2, P1, Pz, P2 |
| Right Parietal (RP) | CP4, CP6, CP8, P4, P6, P8 |
| Occipital (O) | PO3, POZ, PO4, O1, OZ, O2 |

### 2.9. Linear Mixed-Effects Models

As our sample sizes across age groups were unbalanced (DeBruine & Barr, 2021), linear mixed-effect models (LMEs) were implemented using the LME4 and AFEX (Heckerman et al., 2016) libraries for R to test for the effects of Rhythmic Level, Age Group, and Electrode Cluster on neural entrainment to music. Across the LMEs, Satterthwaite’s method was used to estimate degrees of freedom for F-Tests and to compute probability values. Calculating standard effect sizes for LME is an on-going area of research (Jaeger et al., 2017), and not all types of model objects currently have software support for computing effect sizes for mixed models. For models built using the LME4 library, the semipartial (marginal) R-squared was calculated as an effect size for fixed effects (Jaeger et al., 2017) using the developer version of the r2glmm package for R (accessed on GitHub 5/24/2022). For models built using the AFEX library, unstandardized effects (e.g., mean differences) for contrasts of interest are reported using the emmeans library (Lenth, 2022). The global α-level was set to 0.05. In instances of multiple comparisons for the F-Tests that did not involve *a priori* hypotheses (Cramer et al., 2016), Holm’s correction was used to control the family-wise error rate and produce corrected p-values (Holm, 1979).

## 3. Results

### 3.1. Behavioral Battery Results

Older adults did not significantly differ from younger adults in music perception abilities as assessed using the MBEA, Welch’s independent t-test, t(28.932) = 1.458, p = 0.1556, 95% CI = [-0.7806, 4.6556], Cohen’s D = −0.54. However, consistent with previous literature (Belfi et al., 2021; Mas-Herrero et al., 2013; Müllensiefen et al., 2014), older adults scored significantly lower than younger adults in music reward sensitivity as assessed using the BMRQ, Welch’s independent t-test, t(26.028) = 2.4916, p = 0.01942, 95% CI = [1.9869, 20.7131], Cohen’s D = −0.93, and in general musical sophistication as assessed using the Gold-MSI, Welch’s independent t-test, t(28.634) = 3.4111, p = 0.001946, 95% CI = [18.4922, 73.9495], Cohen’s D = −1.27. Table 2 reports the means and standard deviations of age, BMRQ, Gold-MSI, and MBEA in each group.

**Table 2.** Younger and older adults mean values and standard deviations for age (years), the total score of the Barcelona Music Reward Questionnaire (BMRQ), the total score of the Goldsmiths Musical Sophistication Index (Gold-MSI), and scores on the melodic contour perception task from the Montreal Battery of Evaluation of Amusia (MBEA).

|  |  | All Participants | Younger Adults | Older Adults |
| --- | --- | --- | --- | --- |
| Age | Mean | 45.38 | 19.81 | 70.94 |
|  | SD | 26.66 | 1.60 | 8.48 |
| BMRQ | Mean | 75.88 | 80.75 | 71.00 |
|  | SD | 13.85 | 10.75 | 15.17 |
| Gold-MSI | Mean | 184.19 | 203.69 | 164.69 |
|  | SD | 46.06 | 36.79 | 47.12 |
| MBEA | Mean | 23.09 | 23.94 | 22.25 |
|  | SD | 3.74 | 3.73 | 3.68 |

### 3.2. Natural Pulse Frequency of Self-Selected Music Did not Differ between OA and YA

Out of 96 possible musical recordings per age group (16 participants*6 musical recordings for each participant), a total of 77 musical recordings for the YA (N = 16, mean number of recordings per participant = 4.81), and 61 musical recordings for the OA (N = 15, mean number of recordings per participant = 4.07) survived the music-feature analysis and were subjected to the neural-entrainment analysis. To assess the variability of and age-related differences in natural pulse frequencies for self-selected music across the younger and older adults, probability density functions were calculated for the natural pulse frequencies for each age group (Figure 2). Both YA and OA exhibited similar distributions in the natural pulse frequencies of their self-selected music, with a mode in their respective distributions arising at ~2Hz, consistent with previous work suggesting 2 Hz is a frequently occurring pulse frequency in natural music (Ding et al., 2017; Large et al., 2015). A Kolmogorov-Smirnov test of the pulse-frequency distributions suggested that the distributions of the natural pulse frequencies did not significantly differ between the YA and OA groups, D = 0.16, p = 0.36, suggesting that older and younger adults selected music with similar natural pulse frequencies.

**Figure 1.**
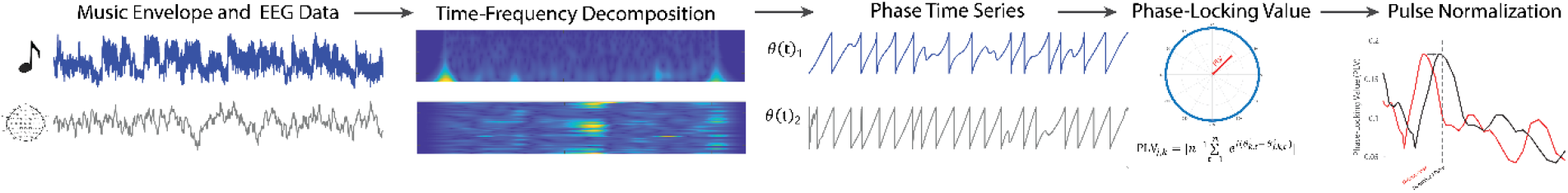
Measuring Neural Entrainment to Musical Pulse. Neural entrainment to musical pulse and rhythm was quantified as the phase-locking value (PLV) between EEG channel activity and the envelope of each musical recording. First, raw EEG channel activity was pre-processed, and the amplitude envelope of each musical recording was estimated. A time-frequency decomposition, using the complex wavelet transform, was then conducted to estimate instantaneous phase information in the EEG timeseries and musical envelopes. PLVs were computed from the resulting phase timeseries. Finally, PLVs underwent pulse normalization, prior to second-level analyses.

**Figure 2.**
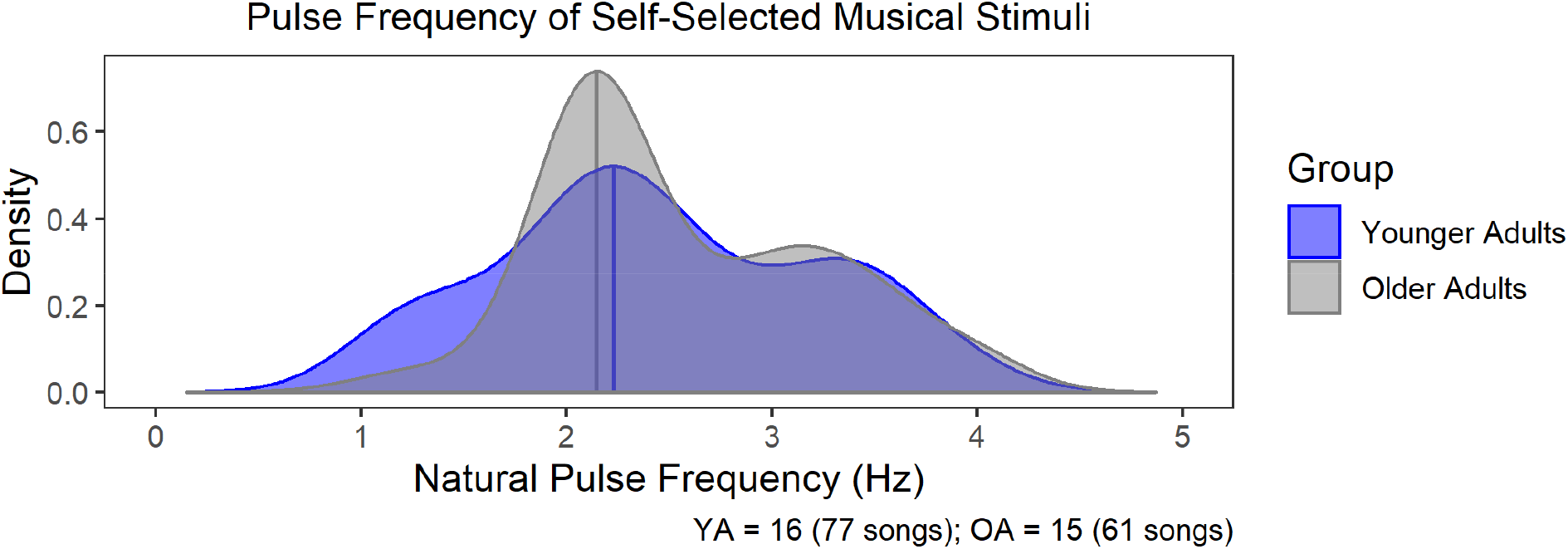
Natural Pulse Frequencies in Participant-Selected Music. Probability Density Functions (PDFs) of natural pulse frequencies for younger (YA; blue) and older (OA; grey) adults’ self-selected musical recordings that survived the music-feature analysis. Younger adults (N = 16, 77 musical recordings) and older adults (N = 15, 61 musical recordings) exhibited similar distributions of natural pulse frequencies in their self-selected music, with a prominent mode emerging ~2 Hz, prior to the pulse normalization of the PLVs. A Kolmogorov-Smirnov test of the pulse-frequency distributions suggested that the distributions of the natural pulse frequencies did not significantly differ between the YA and OA groups.

### 3.3. Neural Entrainment at the Pulse Level Did not Differ between OA and YA

Based on prior research, we predicted *a priori* that, on average, stronger PLVs would emerge at the pulse level, relative to the neighboring off-pulse level (Fiveash et al., 2020). Moreover, we predicted that younger adults would exhibit stronger phase-locking to musical pulse, relative to older adults (Henry et al., 2017). We, first, tested these predictions in a global manner, by using the grand-averaged PLVs for each participant in the YA and OA groups. As shown in Figure 3a, when averaged across all electrodes, younger (N =16) and older adults (N = 15) exhibited strong PLVs at the pulse level, manifested as a local maximum in the PLV plot, consistent with this prediction. Contrary to our predictions, however, the grand-averaged PLVs at the level of the pulse appeared comparable across the YA and OA groups. We implemented a LME model to investigate whether neural entrainment was stronger at the pulse level, relative to the off-pulse level, and statistically different across age groups: in particular, we predicted to observe *a priori* a Rhythm Level*Age Group interaction, reflecting stronger neural entrainment at the pulse level and in younger participants. For the LME, the grand-averaged PLV for each participant was entered as the criterion variable, and Age Group (e.g., YA, OA) and Rhythmic Level (e.g., Pulse, Off-Pulse) were entered as fixed effects with an interaction term. Participants were added as a random effect. As random intercept-and-slope models, with and without correlated intercepts and slopes, failed to converge, participant was ultimately added using a random-intercept model, according to R’s formula notation:

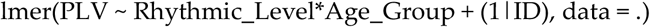

**Figure 3.**
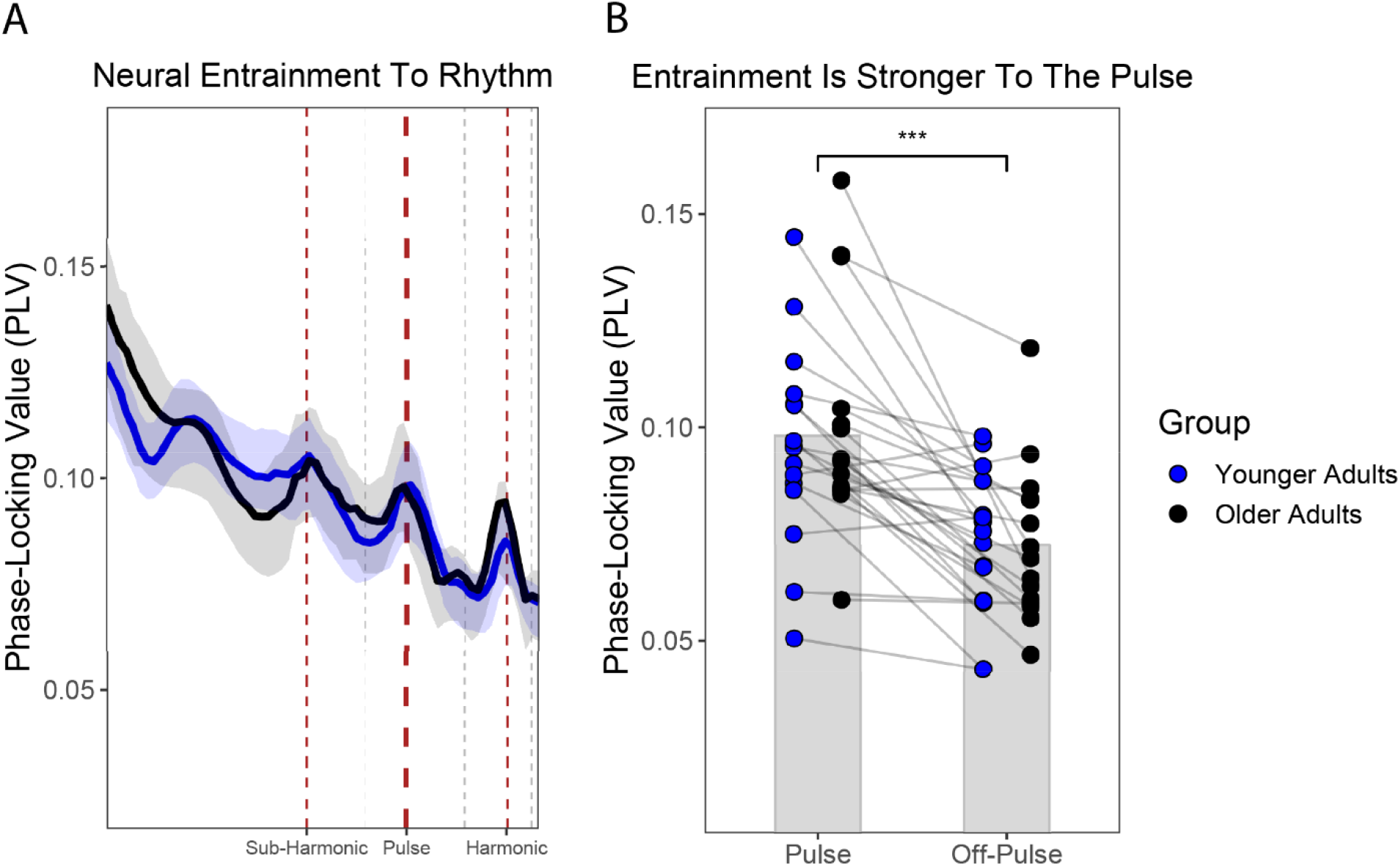
Neural Entrainment to Musical Pulse and Rhythm. A) Phase-locking values (PLVs) grand-averaged across all EEG channels and each participant’s self-selected musical recordings, following pulse normalization of the PLVs. Both younger adults (blue) and older adults (black) exhibited stronger PLVs at rhythm-related levels (e.g., sub-harmonic, pulse, and harmonic levels, respectively; red dashed lines), manifested as local maxima in the PLV plots, compared to off-rhythm levels (e.g., off-sub-harmonic, off-pulse, off-harmonic, respectively; grey dashed lines), manifested as local minima. Shaded errors bars are bootstrapped 95% confidence intervals. B) Grand-averaged PLVs at the pulse and off-pulse levels for each younger adult (blue) and each older adult (black). PLVs were consistently stronger at the pulse level than at the off-pulse level. Grey bars represent the mean PLV for the pulse and off-pulse level, collapsing across younger and older adults (i.e., a main effect of rhythm-level: pulse vs. off-pulse).

As shown in Table 3a, the LME returned a significant main effect of Rhythmic Level, F(1,29) = 35.17, p = <0.001, semi-partial R-squared = 0.144, but not a significant main effect of Age Group, F(1,29) = 0.13, p = 0.72, semi-partial R-squared = 0.005, or Rhythmic Level*Age Group interaction, F(1,29) = 0.21, p = 0.65, semi-partial R-squared = 0.002, suggesting that neural entrainment significantly differed across the pulse and off-pulse levels, but not age groups. A post-hoc test for the main effect of Rhythmic Level (Figure 3b) revealed that the PLVs at the pulse level were significantly stronger, relative to the off-pulse level, t(29) = 5.93, p < 0.0001.

**Table 3a.**
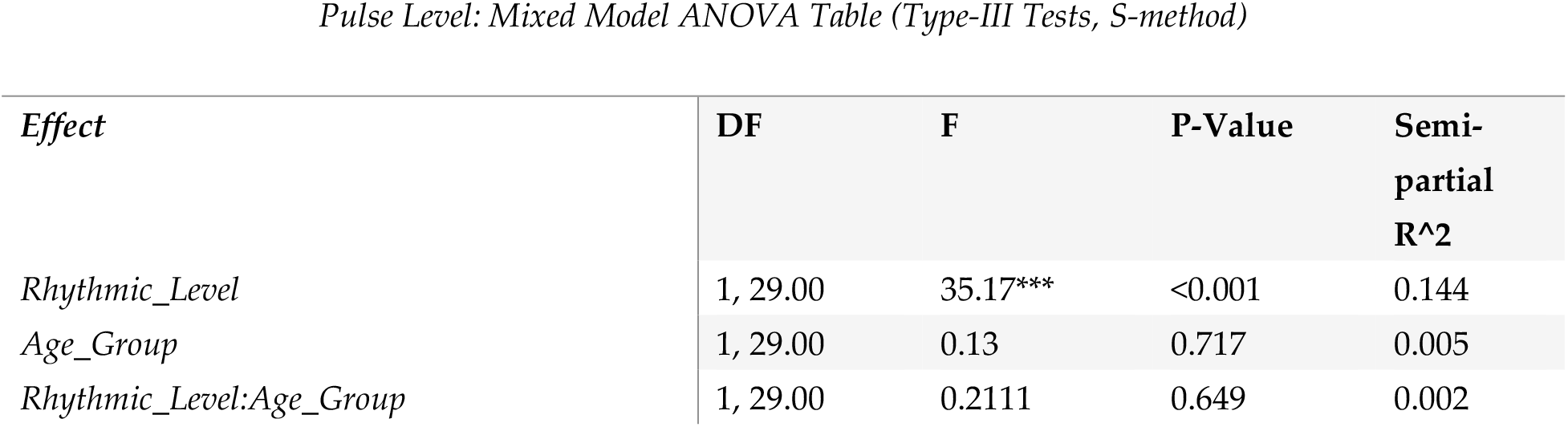
Pulse Level: ANOVA Table. For the pulse level, a linear-mixed effect model returned a significant main effect for Rhythmic Level (e.g., Pulse Level, Off-Pulse Level) on phase-locking values (PLV) to music.

### 3.4. Neural Entrainment to the Pulse Differed Across Electrode Clusters and Age Groups

While neural entrainment to musical pulse did not significantly differ across age groups when the PLVs were grand-averaged across all electrodes, examinations of the topographic representations of PLVs (Figure 4a, Figure 5a,b) indicated there may be age-related differences in the underlying neural networks that are entraining to musical pulse and the sensitivity to the concurrent visual stimulation. For instance, YA exhibited stronger PLVs at the pulse level near a cluster of parietal-occipital electrodes (i.e., near electrode cluster O), relative to the off-pulse level. In contrast, OA exhibited a more uniform topography of PLVs at both pulse and off-pulse levels. We, next, tested whether the topographic representation of PLVs interacted with the pulse and off-pulse levels and the YA and OA age groups. We ran an LME with the PLVs as the criterion variable, Electrode Cluster (e.g., 9 electrodes clusters; defined in Table 1) Age Group (e.g., YA, OA), Rhythmic Level (e.g., Pulse, Off-Pulse) as fixed effects with interaction terms, and participant as a random effect. Because a model with correlated random intercepts and slopes failed to converge, we estimated a LME model using the AFEX library with participants entered as a random effect with uncorrelated random slopes and intercepts--a model which successfully converged:

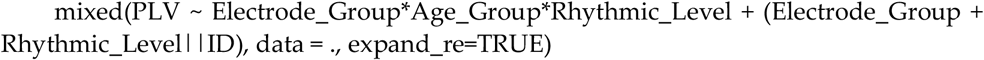

**Figure 4.**
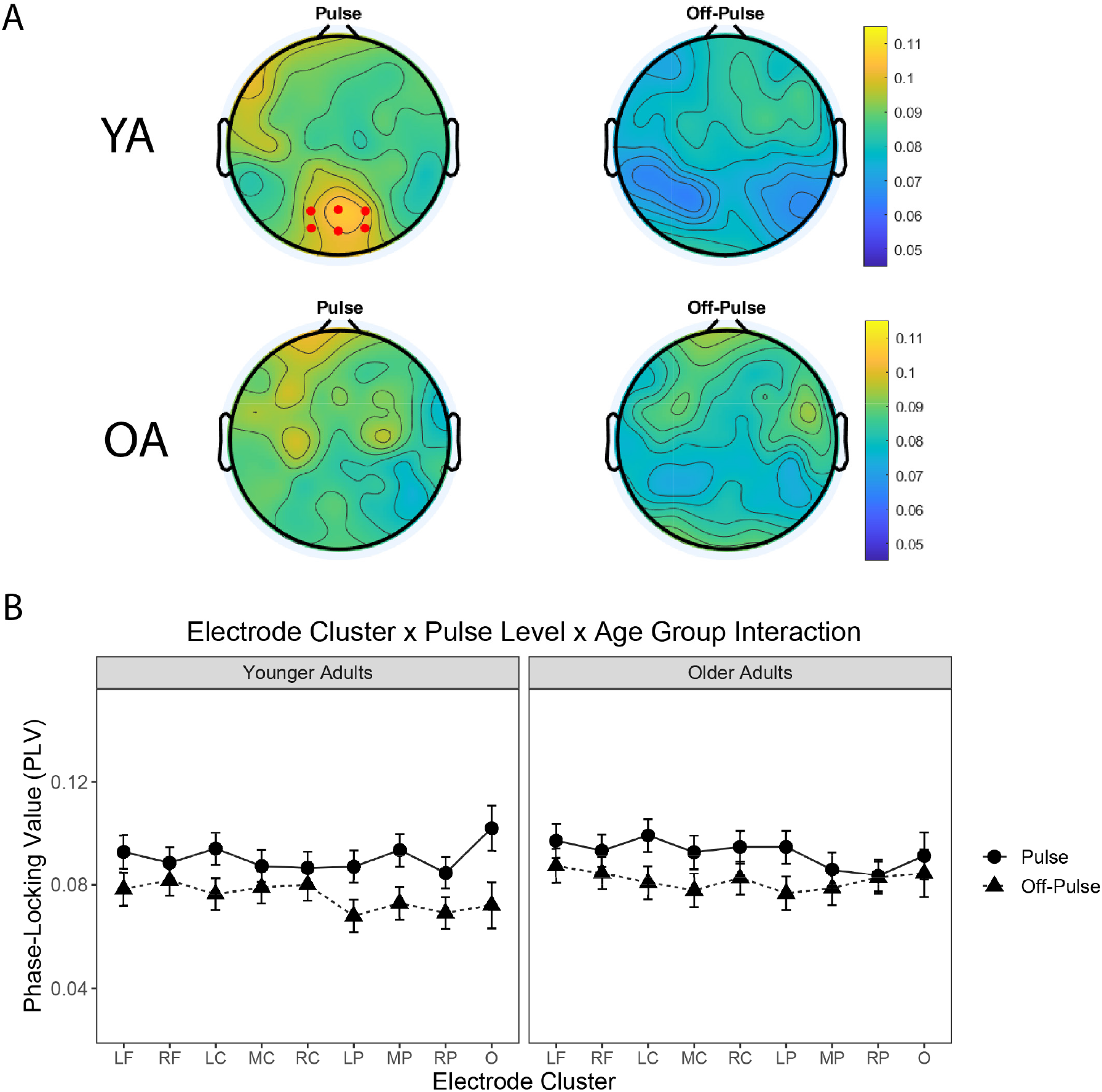
Neural Entrainment Interacts with Electrode Cluster and Age Group. A) Topographies of the phase-locking values (PLVs) for the pulse and off-pulse levels. Younger adults (top row) exhibited higher PLVs at the pulse level in a cluster of parietal-occipital electrodes (e.g., electrode cluster O, red dots denote which channels constitute that cluster). Older adults (bottom row) exhibited a more uniform topography of PLVs at the pulse level. B) Mean PLVs representing a three-way interaction between Rhythm Level (Pulse, Off-Pulse), Electrode Cluster (9 Clusters), and Age Group (Younger Adults, Older Adults). Error bars are standard error of the mean (SE).

**Figure 5.**
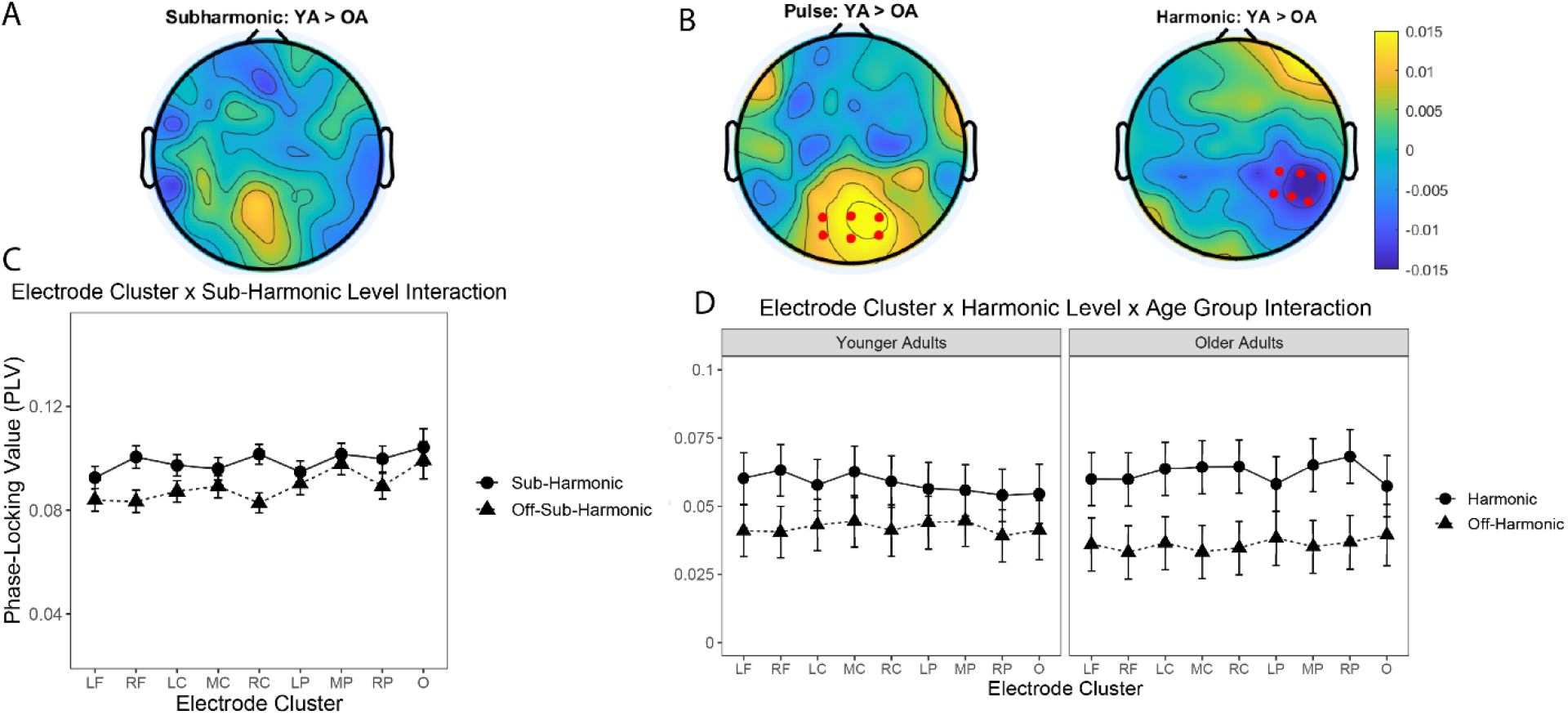
Neural Entrainment to Sub-Harmonic and Harmonic Levels. A) Difference topographies (Younger Adults (YA) > Older Adults (OA) of phase-locking values (PLV) for the sub-harmonic levels. B) Difference topographies (Younger Adults (YA) > Older Adults (OA) of phase-locking values (PLV) for the pulse, and harmonic levels. Red dots on the pulse-level topography represent electrode cluster O, while red dots on the harmonic-level topography represent electrode cluster RP. C) Mean PLVs representing a two-way interaction between Rhythm Level (Sub-Harmonic, Off-Sub-Harmonic) and Electrode Cluster (9 Clusters). Error bars are standard error of the mean (SE). D) Mean PLVs representing a three-way interaction between Rhythm Level (Harmonic, Off-Harmonic), Electrode Cluster (9 Clusters), and Age Group (Younger Adults, Older Adults). Error bars are standard error of the mean (SE).

The LME returned a significant three-way Electrode Group*Rhythmic Level* Age Group interaction (Table 3b), F(8, 3006.57) = 7.53, p < 0.001, p-Holm < 0.001, suggesting that the strength of neural entrainment to music depended on the specific electrode cluster, rhythmic level, and age group (Figure 4b). In particular, younger adults displayed enhanced phase-locking to the pulse level near electrode cluster O. A post-hoc test revealed that younger adults had stronger PLVs to the pulse level, relative to the off-pulse level, at electrode cluster O, t(39) = 4.57, mean difference = 0.0299 (SE = 0.00653). Moreover, the interaction also captured the stronger neural entrainment to the pulse level, in the group of younger adults, at electrode cluster O. A post-hoc test between YA and OA PLVs at electrode cluster O revealed that YA had slightly stronger PLVs at the pulse level, t(191) = 0.842, mean difference = 0.0107 (SE = 0.0128).

**Table 3b.** Pulse Level: ANOVA Table. For the pulse level, a linear-mixed effect model returned a significant three-way interaction between Electrode Group*Rhythmic Level*Age Group fixed effects on phase-locking values (PLV) to music.

| Effect | DF | F | P-Value | Holm P-Value |
| --- | --- | --- | --- | --- |
| Electrode_Group | 8, 57.13 | 2.58 | 0.018 | 0.070 |
| Age_Group | 1, 29.00 | 0.40 | 0.532 | 1.00 |
| Rhythmic_Level | 1, 29.00 | 8.97* | 0.006 | 0.029 |
| Electrode_Group:Age_Group | 8, 57.13 | 0.49 | 0.860 | 1.00 |
| Electrode_Group:Rhythmic_Level | 8, 3006.57 | 5.53*** | < 0.001 | <0.001 |
| Age_Group:Rhythmic_Level | 1, 29.00 | 0.30 | 0.590 | 1.00 |
| Electrode_Group:Age_Group:Rhythmic_Level | 8, 3006.57 | 7.53*** | <0.001 | <0.001 |

### 3.5. Neural Entrainment at Sub-Harmonic and Harmonic Levels Did not Differ between OA and YA

In addition to testing our *a priori* hypotheses for the pulse level, we also conducted exploratory analyses of the neural entrainment to the sub-harmonic (Table 4) and harmonic levels (Table 5) in a series of LME models. Similar to the LME model for the pulse and off-pulse levels, LME models were estimated with electrode Cluster (e.g., 9 electrodes clusters; defined in Table 1) Age Group (e.g., YA, OA), Rhythmic Level (e.g., Sub-Harmonic vs. Off-Sub-Harmonic or Harmonic vs. Off-Harmonic) as fixed effects with interaction terms, and participant as a random effect with uncorrelated random slopes and intercepts. The LME for the sub-harmonic level revealed a significant two-way interaction (Table 4) between Electrode Group and Rhythmic Level (e.g., Sub-Harmonic, Off-Sub-Harmonic), F(8, 3013.38) = 8.23, p < 0.001, p-Holm p < 0.001, suggesting that neural entrainment to the sub-harmonic level was dependent on the specific electrode cluster, but not on age group.

**Table 4.** Sub-Harmonic Level: ANOVA Table. For the subharmonic level, a linear-mixed effect model returned a significant two-way interaction between Electrode Group*Rhythmic Level fixed effects on phase-locking values (PLV) to music.

| Effect | DF | F | P-Value | Holm P-Value |
| --- | --- | --- | --- | --- |
| Electrode_Group | 8, 62.24 | 3.22* | 0.004 | 0.020 |
| Age_Group | 1, 29.00 | 0.16 | 0.693 | 1.00 |
| Rhythmic_Level | 1, 29.00 | 10.33* | 0.003 | 0.019 |
| Electrode_Group:Age_Group | 8, 62.24 | 0.60 | 0.773 | 1.00 |
| Electrode_Group:Rhythmic_Level | 8, 3013.38 | 8.23*** | <0.001 | <0.001 |
| Age_Group:Rhythmic_Level | 1, 29.00 | 0.01 | 0.934 | 1.00 |
| Electrode_Group:Age_Group:Rhythmic_Level | 8, 3013.38 | 1.74 | 0.085 | 0.341 |

**Table 5.**
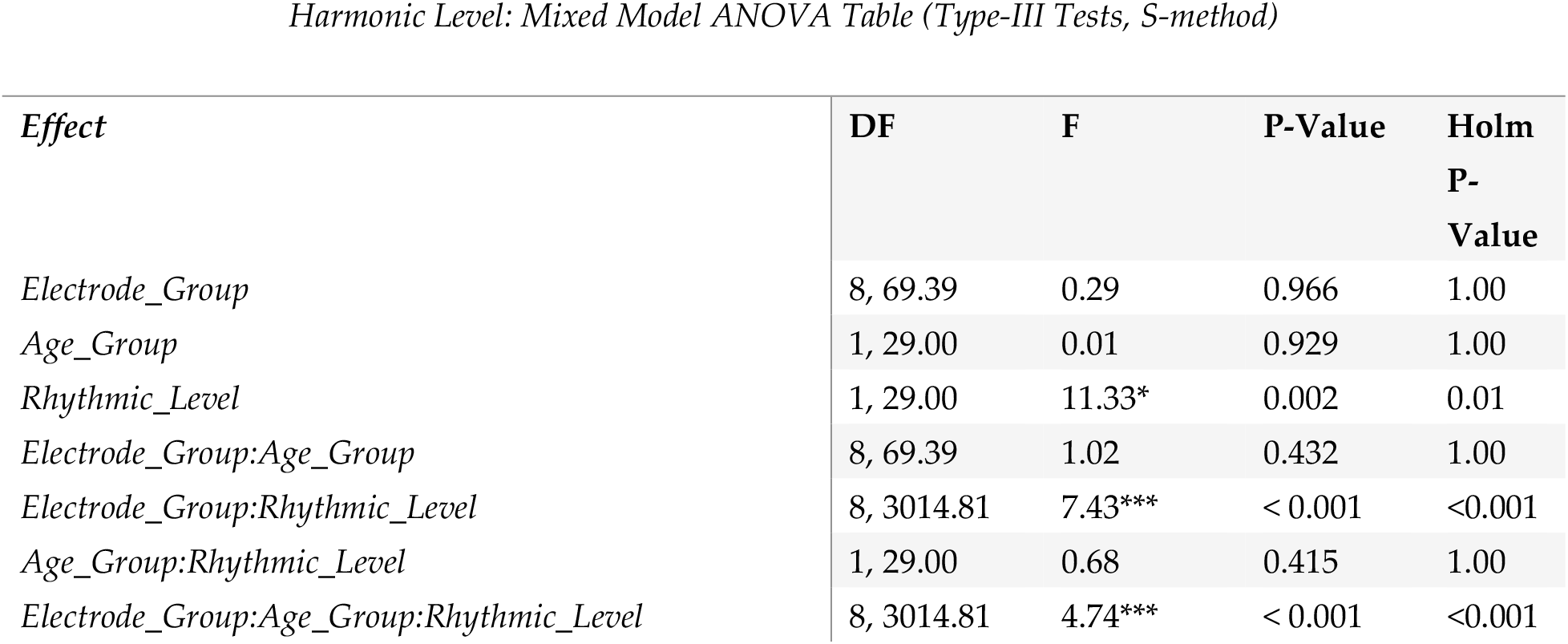
Harmonic Level: ANOVA Table. For the harmonic level, a linear-mixed effect model re-turned a significant three-way interaction between Electrode Group*Rhythmic Level*Age Group fixed effects on phase-locking values (PLV) to music.

Similar to the findings for the pulse level (Table 3), the LME for the harmonic level revealed a significant three-way interaction (Table 5) between Electrode Group, Rhythmic Level (e.g., Sub-Harmonic, Off-Sub-Harmonic), and Age Group, F(8, 3014.81) = 4.74, p < 0.001, p-Holm < 0.001, suggesting that neural entrainment to the harmonic level was dependent on the specific electrode cluster, rhythmic level, and age group. In particular, OA displayed enhanced phase-locking to the harmonic level near electrode cluster RP. A post-hoc test revealed that older adults have stronger PLVs to the harmonic level, relative to the off-harmonic level, at electrode cluster RP, t(31) = 3.41, mean difference = 0.0314 (SE = 0.00653). Moreover, the interaction also captured the stronger neural entrainment to the harmonic level, in the group of older adults, at electrode cluster RP. A post-hoc test between YA and OA PLVs at electrode cluster RP revealed that OA had slightly stronge PLVs at the harmonic level, t(52) = 1.03, mean difference = 0.0142 (SE = 0.0137).

## 4. Discussion

Here, we investigated neural entrainment to musical pulse in self-selected, naturalistic music in a sample of younger adults and older adults. While previous research has demonstrated that the human nervous system entrains to the rhythmic structure of music (Fujioka et al., 2012, 2015; Harding et al., 2019; Nozaradan et al., 2012; Stefanics et al., 2010; Will & Berg, 2007), these studies largely employed artificial auditory rhythms or short excerpts of naturalistic music to study neural entrainment to rhythm, limiting their ecological validity. As part of a larger project on the development of a novel music-based intervention (MBI) for aging, in the current study, participants listened to self-selected music during a period of audiovisual stimulation. Participants were explicitly instructed to select their own music for the audiovisual stimulation, as self-selected music represents a more ecologically valid music listening experience that is linked to efficacy of MBIs (Leggieri et al., 2019) and increases neural activity across auditory and reward systems (Blood & Zatorre, 2001; Loui, 2020; Quinci et al., 2021; Salimpoor et al., 2013).

We analyzed the strength of neural entrainment to musical pulse, quantified as the phase-locking value (PLV) between EEG signals and the amplitude envelope of musical recordings, after normalizing the PLV to the pulse level of each self-selected musical recording following a neurodynamical music-feature analysis. Consistent with our predictions, we observed strong neural entrainment (i.e., relatively higher PLVs) at the pulse level. We also observed neural phase-locking in both younger adults and older adults to other hierarchical levels of meter, specifically at the sub-harmonic and harmonic levels of the pulse. Unlike previous work (Henry et al., 2017), however, the strength of neural entrainment did not reflect a main effect of age at the pulse, sub-harmonic, or harmonic levels, suggesting that neural entrainment to musical pulse and meter may be preserved in aging. Despite no main effect of age, we did observe significant interactions between age and electrode cluster, revealing some age-related differences in neural entrainment at specific channels of EEG activity. In particular, younger adults displayed higher levels of neural entrainment to the pulse level at occipital electrodes, whereas older adults showed stronger neural entrainment to the harmonic level at right temporal electrodes.

While the present study is the first, to our knowledge, to assess neural entrainment to music in a large corpus of naturalistic, participant-selected music across the lifespan, there are several limitations. First, while neural entrainment was measured between electrophysiological signals and the amplitude envelope of recorded music, it is possible that the visual stimulation, during the period of audiovisual stimulation, also shaped neural responses at the pulse and other meter-related levels. Indeed, this may partly explain why younger adults had stronger neural entrainment to rhythm in parietal-occipital electrodes, possibly reflecting additional entrainment to the LED lights from visual cortex, while previous studies with auditory-only stimulation have reported stronger neural entrainment over fronto-central electrodes (Vanden Bosch der Nederlanden et al., 2020; Woods et al., 2021). Secondly, as we permitted participants to select their own music for the audiovisual recording, it is difficult to conclude whether differences in neural entrainment to music across individual participants or across groups (e.g., younger and older adults) are the result of differences in acoustic features of the musical stimuli or differences in endogenous neural function. Nevertheless, our analysis of the natural pulse frequencies for the self-selected music does suggest that the musical stimuli across younger and older adults contained comparable natural pulse frequencies, even prior to the pulse normalization of the PLVs. (In future work, we plan to explore additional recording-specific musical features and acoustic differences (e.g., amount of low-frequency content) into our analyses, as these could contribute to individual and group-level differences in neural entrainment to rhythm (Hove et al., 2014; Weineck et al., 2022).) Acknowledging these limitations of the experimental design, we theorize it is also possible that the topographic differences in neural entrainment to musical pulse could reflect increased sensitivity in younger adults to the visual stimulation that was delivered concurrently with the self-selected music. Indeed, recent EEG results comparing young and older adult groups during light and sound stimulation have shown increased sensitivity in young adults at occipital sites during visual and audiovisual stimulation (Chan et al, 2021), consistent with the idea that young adults are more entrainable than older adults by visual stimulation via lights.

Despite these limitations, this work adds to a growing body of literature that has begun to elucidate possible neurobiological mechanisms underlying the efficacy of MBIs. Recent work, for instance, has demonstrated that the functional connectivity between auditory and reward systems is largely preserved during early stages of dementia (Wang et al., 2020), implicating a possible neurobiological substrate that music-based interventions can target in patients with early-stage dementia. Our findings implicate a complementary neurobiological mechanism—albeit in aging adults without dementia—that music can non-invasively target and entrain rhythmic brain activity in frequency bands that are associated with aging and dementia pathology. To conclude, we believe that assessing neural entrainment to a wider range of naturalistic musical stimuli will help further our understanding how individual differences across expertise, engagement, lifespan development, and various disease states affect musical experiences. Further, we believe it would also illuminate how music can function as a form of non-invasive brain stimulation in designing interventions for healthy aging.

## Author Contributions

Conceptualization, P.T., J.C.K., E.W.L., P.L.; methodology, P.T., E.W.L., P.L.; software, P.T., N.P., J.C.K., E.W.L., P.L.; validation, P.T., N.P., J.C.K., E.W.L., P.L.; formal analysis, P.T., N.P., J.C.K., E.W.L., P.L.; investigation, P.T., N.P., J.C.K., E.W.L., P.L.; resources, P.T., N.P., J.C.K., E.W.L., P.L.; data curation, P.T., N.P., J.C.K., E.W.L., P.L.; writing—original draft preparation, P.T., N.P., J.C.K., E.W.L., P.L.; writing—review and editing, P.T., N.P., J.C.K., E.W.L., P.L.; visualization, P.T., N.P., J.C.K., E.W.L., P.L.; supervision, P.T., N.P., J.C.K., E.W.L., P.L.; project administration, P.T., N.P., J.C.K., E.W.L., P.L.; funding acquisition, P.L., E.W.L. All authors have read and agreed to the published version of the manuscript.

## Institutional Review Board Statement

The study was conducted in accordance with the Declaration of Helsinki, and approved by the Institutional Review Board of Northeastern University (protocol #19-03-20 approved May 14, 2019).

## Informed Consent Statement

Informed consent was obtained from all subjects involved in the study.

## Data Availability Statement

The data presented in this study are available on request from the corresponding author. The data are not publicly available because they are undergoing further analysis as part of a larger study.

## Acknowledgments

We acknowledge MIND Lab research assistants Aaron Kang, Felicia Guo,

